# A microinjection protocol for the greater waxworm moth, *Galleria mellonella*

**DOI:** 10.1101/2024.09.17.613528

**Authors:** James Pearce, Amy Housden, Nicola Senior, Olivia Champion, Joann Prior, Richard Titball, James Wakefield

## Abstract

A limitation to the non-vertebrate 3Rs model *Galleria mellonella* has been the lack of genetic toolkit. A common requirement for genetic tractability is a method to introduce exogenous material to the unicellular embryo, the most common of which is microinjection. This short article describes a detailed method for rearing *Galleria mellonella* to collect large amounts of staged embryos and to dechorionate and microinject embryos with limited mortality.

**Research Highlights:** *Scientific Benefits:* Microinjection allows the introduction of a wide variety of substances, such as DNA, RNA or drugs into *Galleria* embryos, providing the technology needed for genetic engineering, gene editing and functional studies in this important model organism.

*3Rs Benefits:* *Galleria* is being increasingly used as a partial animal replacement model, especially in the field of infection biology. However, uptake has been limited by the lack of genetic and molecular tools. This protocol takes a step towards removing these barriers by providing a means to introduce substances that can create transgenic or genetically engineered *Galleria*.

*Practical Benefits:* Protocol for injecting substances into *Galleria,* using for the most part easily accessible equipment.

*Current Applications:* Generating stable transgenic and gene-edited *Galleria* lines.

*Potential Applications:* Any technique requiring the introduction of substances to *Galleria* embryos. This includes applying existing techniques such as pBac-mediated transgenesis or CRISPR/Cas-based gene-editing to this organism,in order to generate engineered strains of *Galleria*. It could also include injection of synthetic mRNAs encoding proteins fused to fluorescent genes (such as GFP) in order to visualise their dynamics in living embryos; and the injection of drugs that perturb particular cell or developmental processes in order to learn more about early *Galleria* development.

## Introduction

There is a rapidly growing list of publications validating the use of the greater waxworm moth (*Galleria mellonella)* as an *in vivo* animal partial replacement model in the fields of infection, immunology, and inflammation ^1–8^. One advantage of this system is that *Galleria* larvae exhibit an easily identifiable, but qualitative, biological read-out of such challenges – they produce melanin pigment, discolouring the larvae. They possess broad susceptibility to microbial pathogens ^9^, with pharmacodynamics of drug clearance showing remarkably similar patterns of drug clearance to humans^10^. Moreover, individual larvae can be precisely dosed by injection, their maintenance is straightforward and, in contrast with competing non-mammalian systems, such as zebrafish*, C. elegans* and *Drosophila*, they can be reared at 37°C, facilitating research into both host-pathogen interactions at temperatures relevant to human pathogens ^2,11,12^. The annotated genome sequence ^13–16^, along with transcriptomic and proteomic data ^17–27^, provides insight into molecular processes during infection and life history. It also allows development of more advanced methodologies for working with this organism on a macroscopic and molecular level, which in turn can increase uptake and help provide consistent results for researchers who need to progress from a purely *in vitro* environment to in *vivo* testing. Together these advantages present *Galleria* as an eminently suitable partial replacement animal model for rodent systems.

However, the lack of a molecular genetic toolkit with which to engineer *Galleria* for use in animal replacement experiments currently limits its uptake by the research community. The most common method for generating transgenic insects, and other model animals, is to inject a small amount of mutagenic substance into the zygote with microneedles, resulting in the desired modification to the genome. This method of delivery, termed microinjection, is a versatile way of introducing other exogenous substances too, enabling techniques such as RNAi, transient expression of fluorescently tagged proteins of interest or drug delivery.

Insect embryos possess a tough serosal layer known as the chorion, deposited during oogenesis to protect them from desiccation and damage as the develop^28^. This layer can present a challenge to microinjection as the needles must penetrate it to deliver the cargo inside the embryo, often causing damage to either the needle or the embryo. To circumvent this, microinjection protocols for some insect species with hard chorions have used methods that either first introduce a small hole using a tungsten needle ^29^ through which material can be injected or remove the outer chorionic layer entirely ^30–32^. With either of these methods, the resultant embryos are vulnerable to desiccation, unless care is taken to maintain humidity levels in the post injection treatment.

*Galleria* embryos are laid in clusters by the adult female, which use their long ovipositer to deposit them in crevices within a preferred substrate. The individual embryos vary in shape between elliptical and round, measuring around 0.5 mm ^33^ in diameter on their long axis and are glued to both other embryos from the clutch and the substrate by a secretion from the mother. Whilst soft when first laid, the outer chorionic layer quickly hardens and is impenetrable by standard microinjection needles. We therefore sought to develop a protocol that would enable robust separation and dechorionation of early (<2hr old) *Galleria* embryos to enable microinjection of exogenous substances. We also investigated the effects that dechorionation, microinjection and our post injection rearing had on embryonic survival.

## Materials and Methods

### *Galleria* maintenance and husbandry

An inbred “wild type” *Galleria mellonella* colony was started, with the primary purpose of being able to consistently collect large numbers of embryos within short (hourly) time periods. Tru-Larv late-stage larvae (Biosystems Technology Ltd) were fed on an artificial honey diet (Diet 3, Jorjao et al. 2018 ^34^) at 30°C, constant darkness. 50-75 late-stage larvae/pupae were transferred to clear PET sweet jars containing a small amount of diet and allowed to hatch and mate. Embryos were collected for colony maintenance from egg papers – pieces of folded white baking parchment secured with a paper clip at one end – that had been inserted into the adult jars. These embryos were then placed directly onto artificial honey diet and allowed to develop in opaque PET larval jars, with 5um steel mesh between the lid and the jars to reduce L1/L2 larvae escape, until last instar/pupation whereupon they were transferred to PET sweetie jars as above. The number of larval and adult jars put down per week can be varied depending on the number of experimental animals needed and the life stage required, but at least one larval and adult jar per week ensures a regular turn-over and easy ability to expand the colony. This colony has been under continuous culture since 2018 and, since 2022, under the banner of the Galleria Mellonella Research Centre (GMRC), University of Exeter, UK (www.gmrcuk.org).

### Embryo Collection

Adults were kept at 30°C in constant darkness and allowed to lay on egg papers overnight. The egg papers were removed, and the embryos discarded before replacing the clean egg papers into the jars. The moths were then allowed to oviposit undisturbed in darkness before the papers were once again removed and the embryos collected. The age of the embryos was noted as a range, with the oviposition time plus however long they are allowed to develop after. For instance, embryos collected during a 1 hr oviposition window would be labelled as 0-1 hr old.

### Embryo Separation and Dechorionation

In the first 15 minutes post oviposition (PO), the chorion of *Galleria* embryos is soft enough to inject through; however they remain clustered and removal from laying substrate often results in complete embryonic collapse. At 1 hour PO, the chorion has hardened to an extent that attempting to inject through it often results in a needle breakage or requires such a stout needle that the embryo is destroyed.

We therefore experimented with a variety of treatments to try to separate individual embryos from egg clusters and to remove their chorions to allow microinjection (Table 1). During all treatments embryos were agitated vigorously by either shaking at 350rpm or manually with a Pasteur pipette.

**Table 1.** Embryo separation and dechorionation treatments. *Embryonic separation and dechorionation treatments trialled for the purpose of microinjection. During all treatments embryos placed in a petri dish, covered in the solutions and were agitated by either shaking at 350rpm or by manually via aspiration with a Pasteur pipette*.

| Treatment | Embryo Age | Dechoriation Agent Concentration | Dechoriation Agent Time | Effectiveness of Embryo Separation and Dechoriation |
| --- | --- | --- | --- | --- |
| Benzalkonium Chloride | 1-2hrs | 5 % | 90 s | No effect |
| NaOH | 1-2hrs | 0.25M | 5-10 mins | Moderate separation |
| Bleach | 1-2hrs | 5 % active chlorine + 0.05% Triton X-100 | 90s/2 mins | Separated but results in partially dechorionated/collapsed embryos |
| Bleach | 1-2hrs | 2.5 % active chlorine + 0.05% Triton X-100 | 90s/2 mins | Separated but results in partially dechorionated/collapsed embryos |
| Bleach | 0-1hrs | 1.25 % active chlorine + 0.05% Triton X-100 | 2 mins | Separated and fully dechorionated but results in collapsed embryos |
| Bleach + Triton X-100 | 1-2hrs | 1.25 % active chlorine+ 0.05% Triton X-100 | 2 mins | Separated and fully dechorionated |

After dechorionation via bleach, embryos were filtered through a homemade embryo collection basket and rinsed by immersing once in distilled water and then allowing a gentle stream of distilled water to pass over the collection basket filter cloth until the runoff no longer discoloured a piece of blue roll. Embryos were transferred from the filter cloth on the collection basket to glass slides or cover slips using a paint brush cut down to a single hair or a metal dissecting needle. Embryos dechorionated at 0-1hr PO were often too fragile to handle in this fashion. To counter this, embryos were collected after a 1hr oviposition window and allowed to develop for a further hour at 30 °C before dechorionation, which enabled them robust enough for manual manipulation.

### Embryo Microinjection

Three 22 x 22 mm glass cover slips were glued in parallel to a 25 x 75 mm glass slide using nail polish. Dechorionated embryos were then individually aligned on the glass slides, resting against cover slips, using a sterilised paint brush cut down to a single hair and kept in a humidified chamber until injection. Similar to *Bombyx mori*, it is thought that the germ anlage forms on the mid posterior-ventral surface of the *Galleria* embryo. Whilst they do not always have an obvious anterior or posterior pole when dechorionated where this could be deduced (embryos more oval than round), they were aligned so that the long axis ran parallel to the cover slip, and the more flattened surface (if present) parallel to the glass slide.

Injection solutions were then loaded into Eppendorf Femtotip microneedles using a microloader and attached to an Eppendorf FemtoJet + Injectman 4 microinjection system mounted to a Nikon Eclipse TE2000-U inverted microscope. Microneedles were gently opened against the side of a glass cover slip that had been glued to a microscope slide, and embryos were injected with a small amount of injection mix. Injection volume was estimated from the size of droplet (observed as a “clearing” in the region being injected) and we aimed at ∼0.5 nl or a droplet roughly 1/5 diameter of the embryo corresponding to ∼0.8% of the embryo volume.

### Post-injection rearing

After injection, the glass slides holding the embryos were transferred to an inverted petri dish, with 2% agarose in the well of the dish to maintain humidity and a strip of parafilm across the inside of the lid. Embryos were kept at 30 °C for 3-4 days, to allow them to develop until just before hatching.

The *Galleria* diet used contains glycerol and other ingredients that readily absorb atmospheric water and can become very sticky and even liquify if maintained at high humidities for extended periods. Early *Galleria* larvae did not seem to develop well if this occurred and also did not appear to feed well on an agar version of the larval diet (5% agar, 20% larval diet by weight), preferring instead to burrow into it and subsequently die, possibly by drowning.

To maintain high humidity around the embryos and lower humidity in the diet during the initial larval stages, inverted agar petri dishes containing the embryos were placed directly onto normal larval diet in the PET rearing jars with their agar lids still on. Upon hatching, larvae normally crawled across the parafilm on the lid and down to the diet. Embryos were checked at 8 days post injection and any larvae remaining in the dish were transferred to the diet using a single hair of a paint brush. The petri dishes were removed at 12-14 days post injection and embryonic hatch rate calculated but were otherwise left undisturbed. For any subsequent manipulations and genetic crosses, developing G0 larvae can be screened for phenotype once they reach L3/L4 (around 1-1.5cm) and, once pupated, cut out of their pupal casings and crossed with two wild type pupae of the opposite sex. G1 (heterozygous) embryos from these crosses can be screened for phenotype and a stable homozygous stock generated.

### Embryo treatments and Statistical Analysis

The number of embryos used per repeat and the number of repeats for each treatment differed. Batches of between 26-88 (N=732, 15 repeats) were used for untreated embryos, 80-400 (N=2481, 9 repeats) for the dechorionated embryos and 215-290 (N=1076, 4 repeats) for dechorionated and injected. Dechorionated and dechorionated + injected embryos were plated identically, though untreated embryos were placed directly onto parafilm to prevent them moving during transfer. All embryos came from the same parent colony and had the same post treatment rearing.

Data from repeats was pooled and analysed for differences between observed and expected hatch rates using a χ^2^ (Chi-squared) test, with pairwise Z tests with Bonferroni corrections used to determine significances in hatch rate differences between individual treatments.

### Detailed *Galleria* Rearing Protocol

#### Materials for 3 kg Larval Diet

Plastic tub (10 L)

Bench weighing scales (>1500 g, ±1 g)

Weighing boat

Beaker

Large serving spoon

Organic corn meal (750 g)

Yeast extract (450 g)

Organic soy flour (300 g)

Milk powder (300 g)

Organic runny honey (600 g)

Glycerol 99.5% food grade (600 g)

Beeswax pellets (300 g)

Distilled water

#### Materials for Rearing

Larval diet

Opaque white plastic 2 L storage jars for larvae with screw top lid

Clear plastic 4 L sweet jars for pupating/adults with screw top lid

75 micron aperture/35 micron diameter stainless steel woven mesh (Robinson Wire Cloth, SKU S230 x 0.035)

Egg cartons

White translucent wax paper/baking parchment

Paper clips

Scalpel with disposable blades

Blunt tipped tweezers/entomological forceps (Watkins & Doncaster, D4045)

Distilled water

Dual thermo-hygrometer

Scissors/sharp punch

Precision balance

Incubator (optionally with light timer)

Plastic beakers

Plastic tray

Blue roll

Virkon

Disposable plastic waste bags (ideally biodegradable)

### Making Larval Diet

1. Weigh out all the dry ingredients individually and add to the plastic tub and mix manually wearing gloves
2. Weigh and mix the glycerol and honey in a beaker stir to mix
3. Pour the mixed liquid into the dry ingredients and fold to mix. Break up large chunks with your gloved fingers.
4. The diet should not feel very damp or very dry. If it feels too dry, add a little water and mix again with gloved hands - it is normal for the diet to be slightly too wet when first made but will dry out over the next few days as the flour absorbs all the water over the next 24 hrs.
5. Depending on how much diet is needed, the diet can be left in the plastic tub and used immediately or can be stored at room temperature until needed.

### Rearing

#### Adults

1. Place around 50-75 late-stage larvae (5^th^-6^th^ Instar or ∼2 cm long) in clear plastic jar around 30 cm tall along with a small amount of larval diet and label with strain or origin information. A section of fine stainless steel mesh should be placed over the top and secured by a screw top lid. Ensure you poke holes into the lid beforehand, with a sharp punch/scissor tips to allow gas exchange and avoid the build up of excess moisture.
2. Place jar in incubator and label date. This can either be in total darkness or set to a suitable light : dark cycle. 30 °C seems to be the optimal temperature depending on egg mass vs longevity requirements, however a temperature range between 25-37 °C will work. A small beaker of distilled water can be put in the incubator to maintain relative humidity levels between 60-80 % if needed.
3. When the first adult emerges label the jar with the date of first emergence. Then fold a piece of wax paper lengthways so that there are approximately 4-5 folds in it and secure at one end with a paper clip. Place this in the jar so that the bottom end with the paper clip touches the bottom of the jar and the top is secured by the jar lid. This way it can easily be removed without moth escape.
4. Remove paper every couple of days and inspect for eggs. Eggs can be removed by gently scraping them off paper using scalpel blade or with gloved fingers. Egg papers can be reused but should be discarded after signs of mould or excessive faecal spotting.
5. After most adults have died in the jar (normally roughly 2 weeks from first emergence), transfer contents to a plastic bag and freeze both this and the jar for 24 hrs.
6. Discard the bagged waste in a biological waste bin and sterilise the jar and any mesh or apparatus to be reused with virkon, and then leave to air dry.

#### Eggs

1. Eggs can be transferred to a weighing boat and weighed using a precision balance in order to estimate number the number of eggs present or counted/estimated visually.
2. Eggs to be used for stock should be transferred into an opaque plastic jar containing larval diet and labelled with strain or origin information. Roughly 150 to 200 eggs should be placed into around 200-250g amount of diet. Steel mesh should be placed between the top of the jar and the lid. The lid should have ∼10 large holes poked into it and should then be screwed on very tight. Label with that day’s date.
3. The opaque jar should be placed into incubator set at 30°C. A small beaker of distilled water can be put in the incubator to maintain relative humidity levels between 60-80 %, although humidity within larval rearing jars is normally very high due to metabolic production of water.
4. Any eggs not needed for maintenance of stocks can be used for experimental purposes. Those that are not needed should be frozen for ∼24 hours before discarding in biological waste bin.

#### Larvae

1. Larvae should be checked after ∼3 weeks. If they require more food, then this should be added.
2. It is normal for larvae to spin silken tunnels all around the jar that even attach to the lid. However excessive frass build up or any signs of condensation within the jars normally indicate that mould growth is likely to be present or occur imminently. Transfer larvae either to new diet or emergence jars if large enough.
3. When larvae reach around 2 cm/6^th^ instar, empty the larval jars contents into a tray lined with blue roll. Use entomological forceps/blunt tweezers to gently remove larger larvae and transfer to emergence jars in batches of 50-75 (as mentioned in Adults rearing bullet 1).
4. Place in incubator at 30°C
5. Any larvae not needed for maintenance of stocks can be used for experimental purposes. Any larvae and waste diet that is not needed should be transferred a plastic bag and both the jar and bag frozen for 24 hrs.
6. Discard the bagged waste in a biological waste bin and sterilise the jar and any mesh or apparatus to be reused with virkon, and then leave to air dry.

#### Stopping Escapees

1. *Galleria* larvae are prone to wander, especially at first and last instar, unless suitably contained. Steps should therefore be taken to avoid any escape, where larvae could end up in the outside environment or cause damage to their rearing environment (eg. By chewing through wires in incubators).
2. Consider housing the colony within secondary containment within the incubator (e.g. store adult and larval jars in large plastic storage boxes with ventilation holes)
3. All jars should have fine steel metal mesh placed over the opening and be secured on tightly by a screw on lid. Larger larvae are unable to eat through this and the fine diameter means L1 larvae are unable to escape through it. However, very occasionally some L1s will still manage to escape along the threads of the screw top jars.
4. All jars should contain some diet. L1 larvae will naturally wander about until they find food, and this gives them an incentive to stay within the jars rather than explore the wider world. For this reason, it is also beneficial to place eggs directly on the diet.
5. Adult/emergence jars should always contain egg papers once adult moths are present. This provides a nice oviposition surface and reduces the amount that they lay elsewhere in the jar. Equally egg papers should be cleaned of eggs at least twice a week to prevent L1s hatching within the jars.
6. To monitor levels of L1 escapees, keep small petri dishes of diet in the incubator. These act as bait traps to mop up any especially adventurous larvae, but also allows you to survey for any larger scale jailbreaks.
7. To prevent cross contamination between strains it is worth considering separate incubators and/or allowing embryos to hatch in flugged fly vials to prevent L1 larval ingress/egress and transferring to screw top jars when they are larger.

### Detailed Microinjection Protocol

#### Materials

Blue roll (Kim Wypes)

Petri Dishes

Dechorionation basket (50 ml falcon tube, conical bottom cut off and with the centre of the lid cut out)

Glass beaker

Plastic Pasteur pipettes

Nylon mesh

Plastic squeezy bottle

25×75 mm glass microscope slides

22×22 mm glass cover slips

Clear nail varnish

70% ethanol

Embryo manipulator (dissection needle, paintbrush cut down to single hair etc.)

Parafilm

Injection needles (Femtotips or hand pulled to similar dimensions as those used for drosophila)

#### Preparation of reagents

Dechorionation medium: 50 ml Household thin bleach (4.5% active chlorine), 150 ml ultrapure water, 100 μl Triton X-100 – made no more than 1 week before use

Embryo Wash: 0.05 % Triton X-100 solution – store in plastic squeezy bottle to rinse surfaces

Injection Buffer: 5 mM KCl, 5mM Phosphate buffer pH 7.4 – filter sterilised and frozen in 1 ml aliquots. Spin at 12000 g for 1 min before use. Alternatively, 10x injection buffer can be made and stored.

Agar plates: 2 % Hi-Res agarose solution is poured into petri dishes – made on the day

#### Embryo Collection

1. Cut a 20 cm x 30 cm piece of parchment paper and fold along the long axis back on itself in 2 cm folds to form an accordion shape. Secure at one end with a paper clip.
2. Insert into jars containing adult moth jars, keeping the moths in the light for as short a time as possible before returning the jars to 30 °C and constant darkness for 1 hr.
3. Remove egg papers after one hour and collect embryos by gently brushing down the unfurled paper with gloved hands with some blue roll beneath. The cream embryos can be hard to see sometimes on white baking parchment so holding to the light can help you spot any embryos you may have missed.

#### Dechorionation

1. Collect embryos in a petri dish and incubate at 30 °C in darkness for one hour.
2. Prepare the dechorionation baskets by securing a 5 cm x 5 cm piece of nylon mesh between the cut out lid and the now open tube, and fill the glass beaker with water
3. Cover the embryos with dechorionation solution and incubate with agitation from a Pasteur pipette for 2 mins
4. Decant the dechorionation solution into the dechorionation basket and rinse any remaining embryos from the petri dish using the embryo wash
5. Dunk the basket in the beaker of water for 10-20 s. The embryos might initially float but should sink quickly due to the residual Triton.
6. Remove the dechorionation basket from the beaker and rinse the external sides of the basket under a stream of ultrapure water, allowing the water to flow over the bottom of the mesh. This removes remaining bleach but does not cause damage to the embryos from a direct stream of water nor does it disturb the embryos causing them to stick to the plastic falcon tube.
7. Place the dechorionation basket on some blue roll. When both no bleach can be smelled, and the blue roll does not discolour pink then the embryos have been sufficiently rinsed.

#### Microinjection

1. Prepare slides to mount embryos by using a small blob of nail varnish to glue three 22 x 22mm cover slips in a row down the centre of a glass microscope slide. Clean with 70 % ethanol and leave to dry.
2. Sterilize a micromanipulator and use it to align freshly dechorionated in parallel on the microscope slide, resting against the raised lip formed from the cover slips. Around 300 should fit on one slide using both sides of the cover slips. Do not use embryos that are still partially chorionated or that easily collapse upon gentle manipulation as these will cause needle breakage or collapse upon injection.
3. Cut a strip of parafilm and press into lid of agar plates to secure, which helps stop the slides moving around and provides an easier surface for the L1 larvae to walk on. Invert plate and store embryo slides in the agar plates attached to parafilm to prevent desiccation before injection
4. Load needle with you injection solution (injection buffer + your desired cargo). It is advisable to spin injection solutions before use (1 min at 12,000 x g) to pellet any DNA or protein aggregates that may block the needle and to transfer the top 90 % of the solution to a fresh tube.
5. If you can adjust the angle of your microinjector, use a needle angle of less than 30° from horizontal to enable the needle to enter the embryo more easily
6. Start with pressures of ∼350 h:100 hPa for injection and compensation respectively on your microinjector. Both will need to be adjusted to lower pressures if the needle aperture becomes wider.
7. Open the needle, breaking a small bit of the tip by gently pressing it against the side of the glass cover slips on the injection slide, where a gap can be found between embryos.
8. Using a micromanipulator and pump, inject each embryo with ∼0.5 nl, which corresponds to a droplet roughly a fifth of the diameter of the embryo diameter. Droplets will be observed as a “clearing” within the injected embryo, and you should see it plump up slightly. If high embryo pressure causes them to burst or injection mix to spill out after injection, consider desiccating the embryos by leaving out at room temperature for 5-10 mins.

#### Post Injection Rearing

1. Post injection, store embryos in the inverted agar plates with parafilm for 3-4 days at 30 °C in darkness, before putting the plates into larval rearing tubs containing diet. Do not remove the dish containing the agar or the embryos may desiccate and fail to hatch. They are able to crawl through the small vents in the petri dish instead to reach the diet.
2. Hatch rates can be recorded after 10-14 days by observing the number of unhatched embryos left on the slides, since dechorionated embryos that hatch will leave little trace on the slide.

## Results

Early attempts to inject untreated embryos often failed due to needle breakage against the hard chorionic layer. Trials were therefore undertaken to soften or remove the chorions of young (1-2 hr) embryos. Exposure to NaOH or benzalkonium chloride solutions, as suggested in the literature for other lepidopterans ^35,36^,at various concentrations had little effect. Protocols for dechorionation of *Drosophila* embryos using 2.5-5 % sodium hypochlorite solutions for 1.5-2 mins were able to dechorionate and separate embryos but seemed too aggressive. They resulted in embryos that were damaged on sides fully exposed to the hyperchlorite solution but that remained attached to their neighbours or still partially covered by their chorions. A weaker solution of 1.25% (v/v) hypochlorite for 2 mins was therefore trialled, very similar to that used by Cosi et al. and Abidallah & Roversi ^37,38^, which resulted in embryos that were both fully separated and the majority completely dechorionated. We added 0.05% (v/v) Triton X-100 (commonly used in *Drosophila* embryo wash) to all the bleach solutions trialled, with the primary purpose of preventing embryos floating in the surface film or sticking to plastic surfaces. Cosi et al. ^37^ reported that addition of another non-ionic surfactant, Tween 80, had a protective effect against mortality induced by chlorox treatments; however, we did not investigate the protective effect of low concentrations of Triton X-100.

Having devised a method that reliably allowed us to obtain dechorionated embryos that were <2 hrs old, we investigated the effects on survival that our protocol for dechorionation, handling and subsequent rearing procedures had on their viability. In addition to this we investigated how hatch rate success was impacted by injection with moderate amounts of injection buffer. There were significant differences in hatch rates between both untreated embryos (87 %) and dechorionated (77 %) ones (*P* < 0.05) and untreated and injected (69 %) (*P* < 0.05), as well as between those dechorionated and those which were injected (*P* < 0.05). Although not quantified, embryos from all treatments developed normally post hatching and were able to produce fertile offspring.

Mould was occasionally seen growing on the remaining unhatched embryos after 14 days when calculating hatch rates. This did not appear to affect embryo viability and sterilising injection slides and embryo manipulators before use helped to reduce this. However, from early experiments, we found excess humidity to be a significant issue, causing L1 larvae to drown and die. We prevented this by removing any condensation from the plates before use, and opening plates briefly if they need to be moved from 30°C to room temperature.

## Discussion

This protocol describes, in detail, a method to dechorionate, inject and rear early *Galleria* embryos to adulthood in the laboratory. We show that both dechorionation and injection result in a slight but significant loss in embryo viability at this early stage of development, with roughly 70 % of eggs expected to hatch, provided the injection solution is not in itself toxic. This presents a good starting point for generating new transgenic strains, especially given some insect species have similar hatch rates to this for untreated embryos ^39^.

Our experiments are also consistent with earlier findings ^37,38,40^ that a low strength chlorox solution is suitable for chorion removal in older (24hr) *Galleria* embryos. Whilst we did not investigate the protective effects that the addition of low concentration of non-ionic surfactant added to dechorionation solution has on embryonic survival, we did observe that the age of the embryo at time of treatment has an effect. 0-1 hr embryos are unsuitable for dechorionation with 1.25% chlorox solutions since this renders them so fragile that they collapse under their own weight or at the slightest touch. This may be important for future transgenic techniques where time of injection can affect the efficiency of transformation.

This protocol will hopefully be accessible to many laboratories worldwide. Aside from the microinjection equipment – which is similar in set up to many *Drosophila* and zebrafish apparatus – all the materials are cheap and most readily available within a laboratory environment. The protocol presented is simple and due to the high fecundity of this moth, a single researcher can prepare and inject from 600-800 embryos at a time in a 4-5 hr window. In addition, *Galleria* colonies are easy to maintain with diet ingredients being cheap when bought in bulk and only an incubator required for housing (rearing is possible at lower temperatures but development is slower). A single colony can produce large numbers of all life stages, which may be important for researchers who wish to use larvae for assays at the same time. The low monetary and time cost for researchers should be an advantage for those looking to start using this insect model.

In summary, we have developed a methodology to reliably inject *Galleria* with solutions, whilst maintaining viability through to adulthood. This technique is, in principle, applicable to solutions containing small molecules, proteins or nucleic acids. As such, it is potentially suitable for injection of synthetic mRNAs encoding proteins fused to fluorescent genes (such as GFP) in order to visualise their dynamics in living embryos; and the injection of drugs that perturb particular cell or developmental processes in order to learn more about early *Galleria* development. More importantly, perhaps, it paves the way for attempts at pBac-mediated transgenesis and CRISPR/Cas-based gene-editing, in order to generate engineered strains of *Galleria*. Such strains could transform the use of *Galleria* as a partial mammalian replacement model in the fields of infection, immunology and beyond.

**Fig 1.**
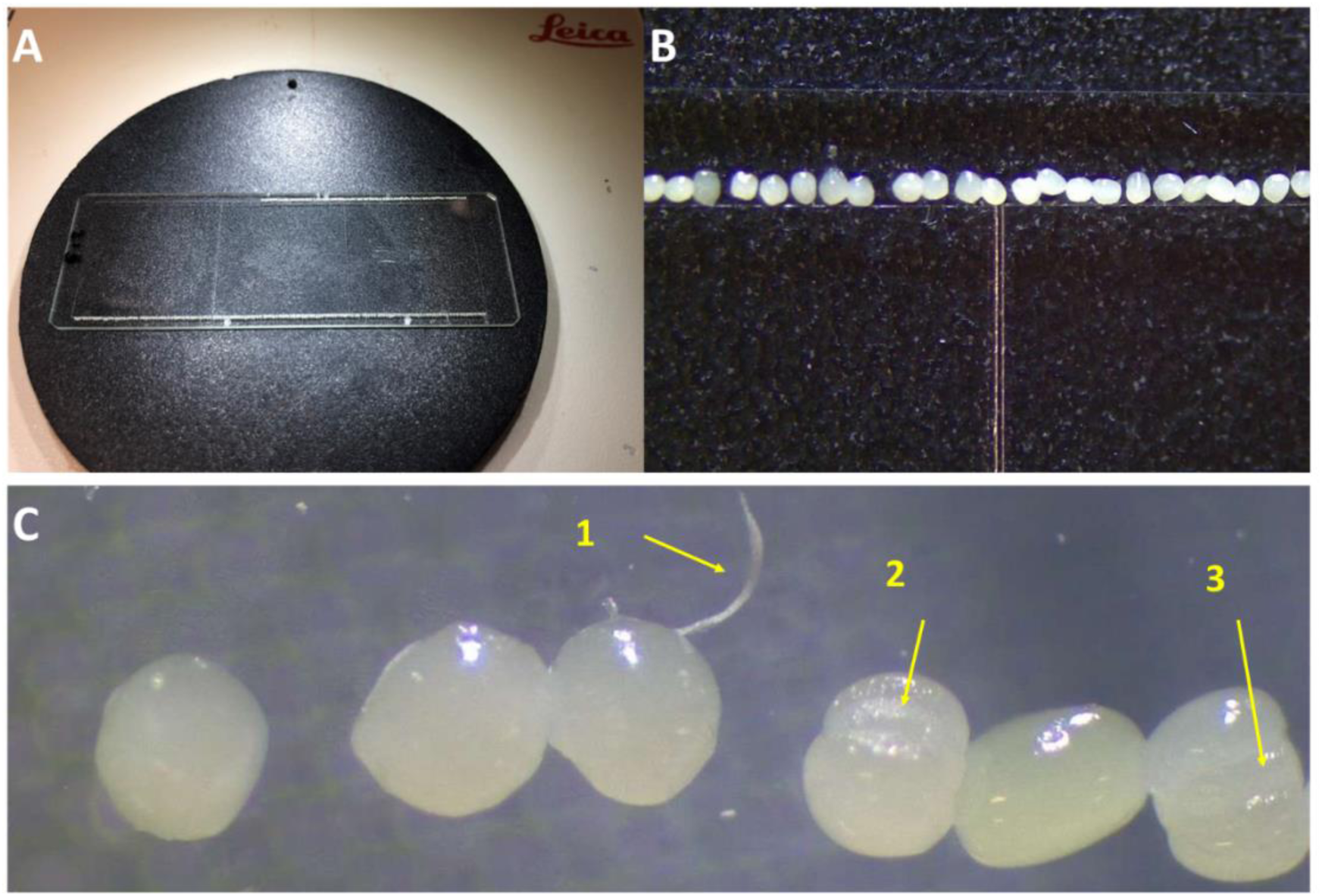
Dechorionated embryos – good vs bad and alignment on cover slip/slide. Embryos can be aligned for injection on glass slides along the edge of cover slips glued down the centre (Image A and B). The cover slips help prevent embryos sliding during injection. Embryos that remain partially chorionated (C - yellow arrows) which present as “wings” (1) or slightly dumbbell shaped (2,3) can cause needle breakage during injection so should not be used.

**Fig. 2.**
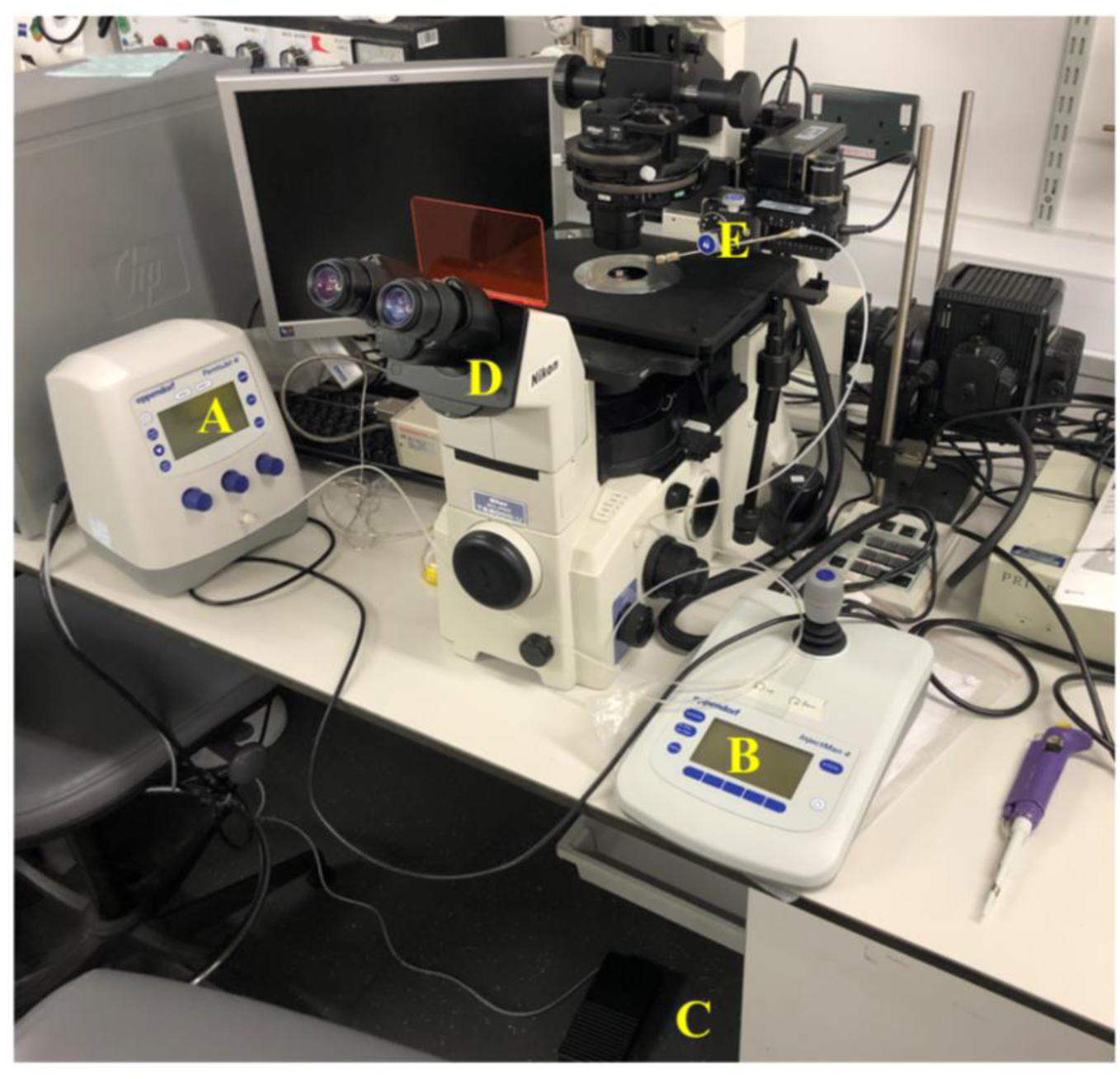
Microinjection Set Up. Galleria *injection set up. An Eppendorf FemtoJet (A) and InjectMan 4 (B) system with foot pedal (C) is mounted to a Nikon TE-2000U inverted microscope (D) enabling precise manipulation of the needle holding apparatus (E) and injection volumes*.

**Fig. 3.**
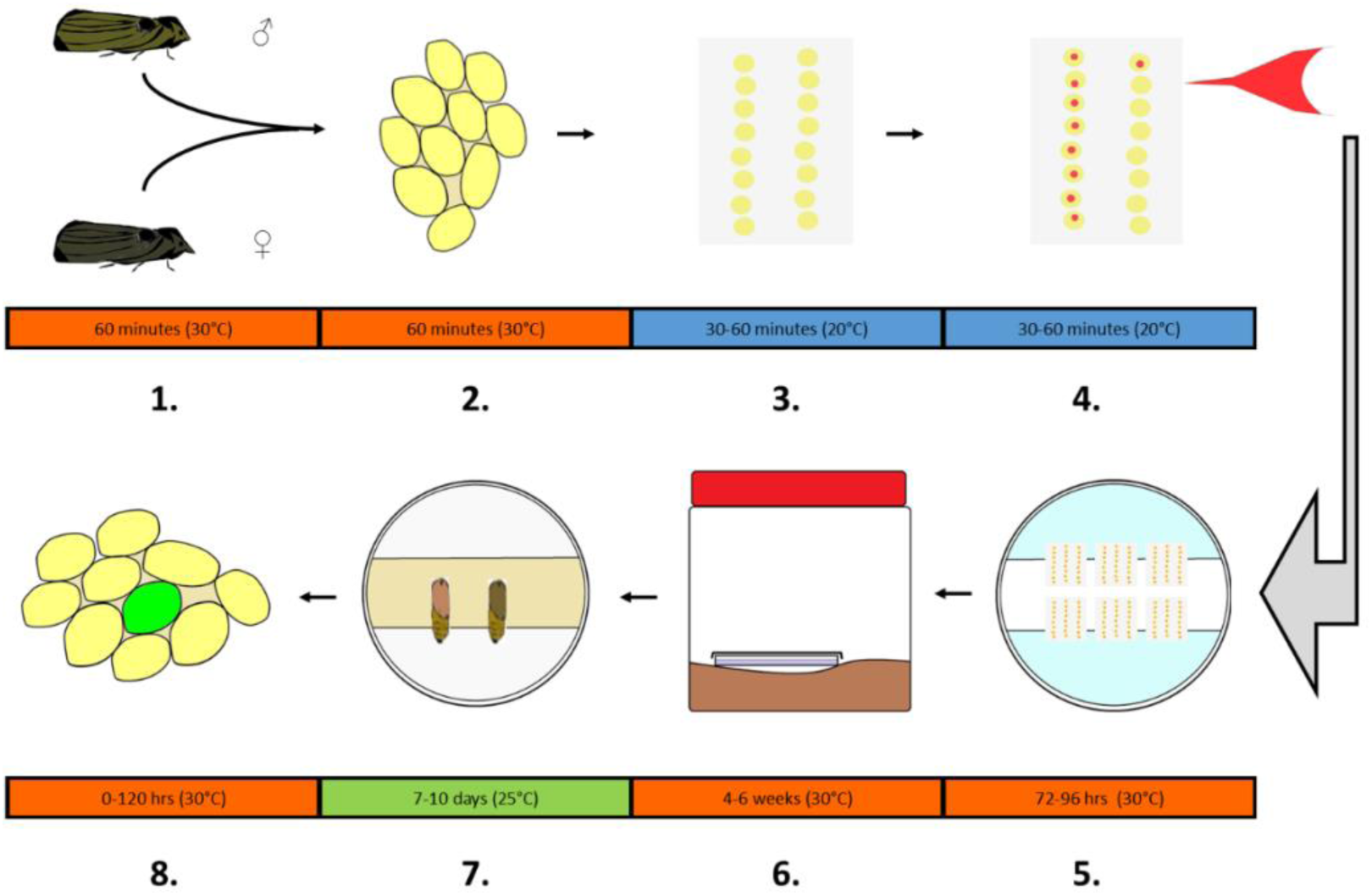
Microinjection Pipeline. Galleria *injection and screening pipeline: **1.** Eggs are obtained from mixed sex adult* Galleria. ***2.** Embryos are left to develop to desired stage. **3.** Embryos are aligned on coverslip for microinjection. **4.** A small amount of microinjection mix is injected into the desired part of the embryo. **5.** Injected embryos are allowed to develop at high humidity. **6.** The embryos are moved to a jar containing larval diet, and the larvae are left to develop undisturbed until they reach late larval stage. **7.** Late larvae can be screened for phenotype or at the onset of pupation, pupae are removed from the sealed container and removed from their casings to allow sexual identification. Individual crosses between virgin injected and wild type adults can then be performed in petri dishes. **8.** Embryos collected from these crosses for screening*

**Table 1.**
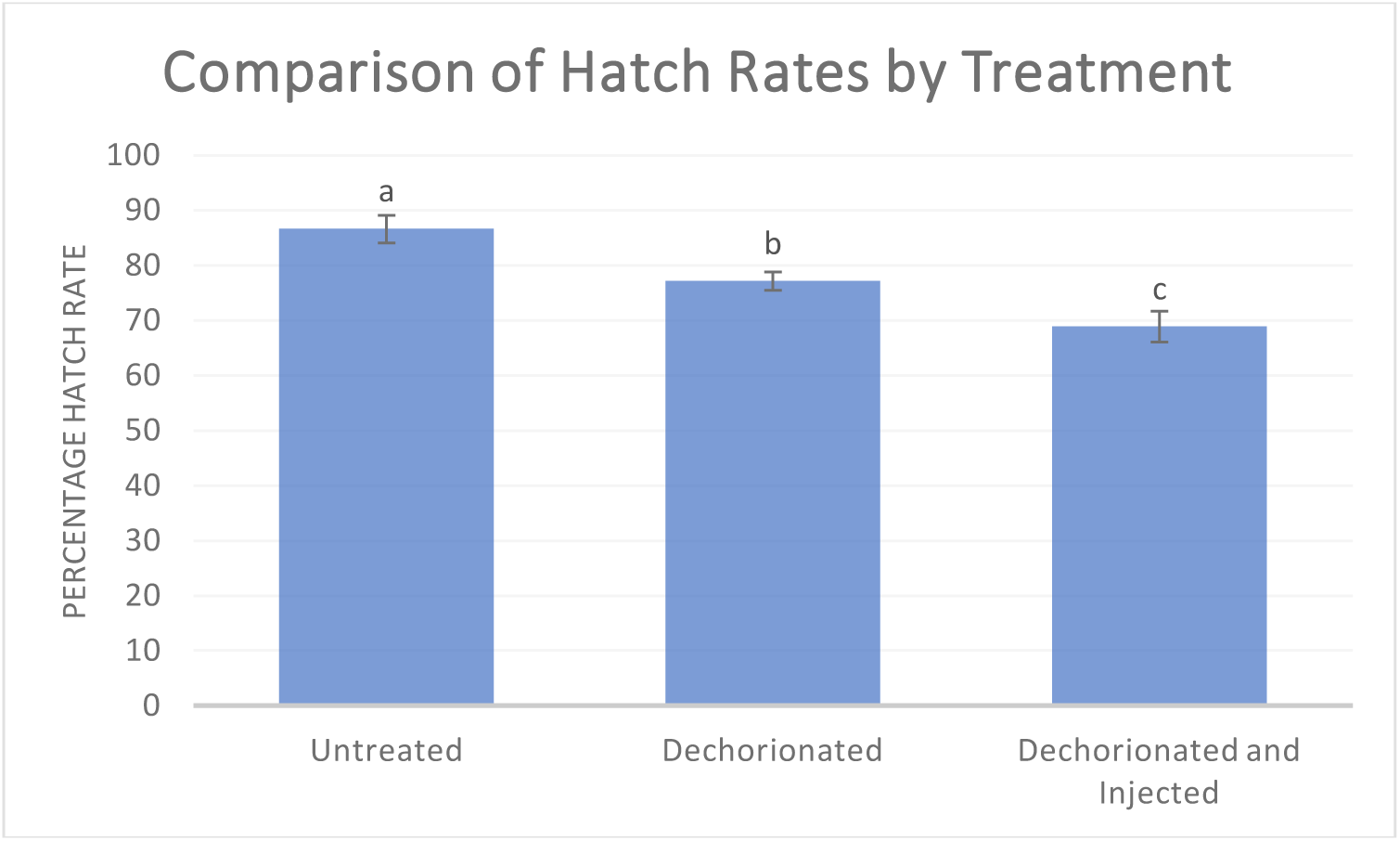
Hatch rates and survival to pupation for different treatments. *Hatch rates of untreated wild type embryos, and those that have been dechorionated or dechorionated and then injected. Different letters above bars indicate significant difference between treatments (P < 0.05, Pairwise Z-tests with corrections). All embryos were subjected to the same post treatment rearing. Error bars represent 95% confidence intervals*.

